# Associations between structural and functional hemodynamics in the optic nerve head

**DOI:** 10.64898/2026.11.17.767382

**Authors:** Yuankai Lu, Yi Hua, Ruhani Gill, Andrew Theophanous, Po-Yi Lee, Ian A. Sigal

## Abstract

**Purpose:** Glaucoma is characterized by progressive degeneration of retinal ganglion cells axons thought to initiate within the optic nerve head (ONH). Although the precise mechanisms remain unclear, a leading hypothesis suggests that insufficient blood flow and oxygenation may be significant contributors to axonal damage. Measuring ONH oxygenation directly, however, is difficult. Hence, our first goal was to identify associations between structural and functional hemodynamics parameters (flow and oxygenation). We then evaluated whether local tissue oxygenation and hypoxia can be estimated from measurable parameters.

**Methods:** Following previously reported methods, the ONH vasculature of four healthy monkey eyes was reconstructed. Numerical simulations were performed to determine blood flow and oxygen distribution. Structural parameters of each segment, including spatial location, orientation, length, and tortuosity, and functional hemodynamics parameters of each segment, including blood flow, pressure, blood oxygenation, and surrounding tissue oxygenation, were quantified.

**Results:** Structural parameters exhibited no distinct spatial distribution, whereas hemodynamic and oxygenation parameters followed radial patterns. Although blood flow correlated with tissue oxygenation (p<0.001), regional mismatches indicated that flow alone was insufficient to predict accurately local oxygenation. We developed two multivariable mixed-effects models using measurable structural and flow parameters, a quadratic model that predicted continuous tissue oxygenation with high accuracy (R^2^=0.75) and a logistic model that estimated local hypoxia risk.

**Conclusions:** Our results provide a framework for estimating local tissue oxygenation and hypoxia risk that currently cannot be measured experimentally from vascular structural and flow parameters that can be measured.

## 1. Introduction

Glaucoma, a leading cause of irreversible blindness worldwide, is characterized by the progressive degeneration of retinal ganglion cells and their axons.^1,2^ This damage is thought to begin in the lamina cribrosa (LC) region^2^, located at the back of the eye within the optic nerve head (ONH). Although the precise mechanisms remain unclear, one leading hypothesis suggests that insufficient blood flow and oxygenation within the LC may be a significant contributor to axonal damage.^3–11^

The LC is a specialized structure crucial for maintaining the health of retinal ganglion cell axons where they exit the globe. At the LC structural support is provided collagenous beams, whereas oxygen and nutrients are provided by a vascular network^5,12–14^ From fundamental fluid mechanics and experience in other tissue^15,16^, it is expected that, in the LC, the functional aspects of the hemodynamics – blood and oxygen supply, depend strongly on the structural characteristics of the LC vasculature – vessel tortuosity, length, orientation, etc. In other words, associations between LC structural and functional hemodynamics parameters are expected.

Parameters that can capture hemodynamics structure and function have been developed for the whole vascular system of entire organs, such as cluster coefficients, connectivity and transport energy consumption.^17–19^ These measures, while useful, are not ideal for the question of glaucoma susceptibility because they do not capture local blood flow and oxygen supply that are of great interest in understanding the ONH and the regional variations that could help explain local neural tissue damage^20^.

Several imaging techniques have been developed to assess structural and/or functional aspects of ocular hemodynamics, including laser speckle flowgraphy (LSFG)^21^, optical coherence tomography angiography (OCT-A)^22–25^, ultrasound techniques^26,27^, and MRI-based techniques^28^. While these have generally been successful for assessing retinal hemodynamics, they are inadequate for the ONH hemodynamics, mainly due to limitations in signal depth or resolution (both spatial and temporal). Given these limitations, theoretical and numerical models have been used to investigate local ONH hemodynamics; however, many rely on simplified vascular geometry and are constrained by high computational costs and the structure – function relationship in ONH hemodynamics remains unknown.^7,29–37^ Recently, we developed techniques based on histological sections to reconstruct 3D ONH vasculature structure.^38^ Combining 3D eye-specific ONH vasculature and high-efficiency hemodynamic simulation, we can derive a detailed 3D map of ONH vasculature architecture and the corresponding hemodynamics and oxygenation and evaluate their relationships in a reasonable time.^39–42^

Our primary goal was to characterize several structural and functional hemodynamics parameters of the monkey ONH, and to identify associations between the parameters. To achieve this objective, four eye-specific 3D models of the ONH vasculature were reconstructed following the approach previously described^38^. We used numerical simulations to determine ONH hemodynamics and oxygenation in each of the models. We then analyzed ONH structural and functional hemodynamics parameters and evaluated their statistical associations. It is important to emphasize that our “functional” parameters are so only from the perspective of hemodynamics and flow and that these should not be confused with functional parameters in terms of vision or biological activity in the eye.

Although tissue oxygenation is of great interest for assessing local neural tissue health and potentially susceptibility to damage, it remains difficult to measure directly in the ONH. Other structural and flow parameters, however, are more accessible experimentally. Hence, a secondary goal was to determine whether local tissue oxygenation could be inferred from the more readily measurable vascular features. Specifically, we developed two multivariate mixed-effects models using these predictors: a regression model for tissue oxygenation, and a logistic model for the probability of local hypoxia, evaluated using a Precision-Recall curve.

## 2. Methods

*General workflow process.* Four eye-specific ONH vasculatures of healthy monkeys were reconstructed using the procedure described before^38^. Briefly, the vasculatures were perfusion-labeled, then the tissues were fixed and sectioned using histology techniques. The sections were imaged and the vessels identified and reconstructed using image segmentation, skeletonization and 3D model reconstruction techniques (Figure 1). We then used the reconstructed vessel networks to do numerical simulations and evaluate the blood flow and oxygen distribution within the ONHs.

**Figure 1:**
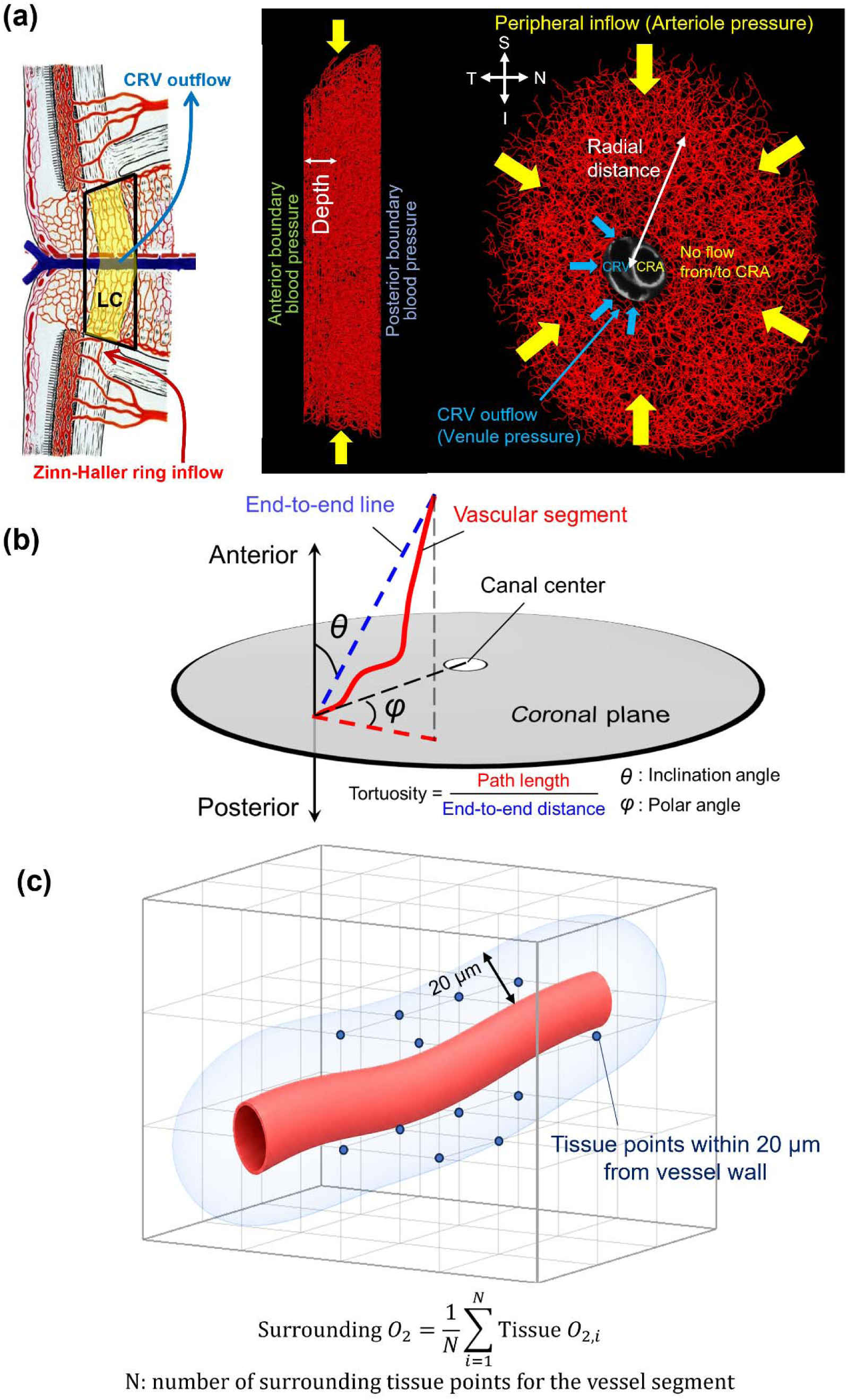
(a) Left: Schematic illustration of the sagittal view of ONH vasculature, adapted from^41^ The central yellow region represents the LC region. The modeled ONH region includes vessels within the scleral canal, bounded peripherally by the connective tissues of the sclera and/or pia mater and centrally by the central retinal artery and vein. The anterior and posterior boundaries are defined by planes perpendicular to the central retinal artery and vein axis, positioned to ensure that the modeled region completely encloses the LC. Blood enters the ONH via the circle of Zinn-Haller (red arrows) and drains through the central retinal vein (CRV, blue arrow). The trapezoidal region outlined within the ONH indicates the modeled region. Right: Two orthogonal views of the reconstructed vascular network. The sagittal view corresponds to the trapezoidal region on the left, depth is defined from the anterior boundary. The coronal view shows the inflow at the periphery (yellow arrows) and CRV outflow (blue arrows). The central retinal artery (CRA) is present but has no flow exchange with the vasculature. Peripheral distance was defined from the ONH canal. (b) Diagram illustrating the definition of the four structural parameters of a vessel segment: length, inclination angle (θ), polar angle (φ), and tortuosity, adapted from^38^. (c) Diagram illustrating surrounding tissue oxygenation for a vessel segment, defined as the average oxygen partial pressure of tissue points within 20 µm of the vessel wall. These are explained in detail in the main body of the manuscript.

We quantified five structural parameters for each whole ONH vascular reconstruction: segment number, branch point number, ONH volume, total vessel length, and average nearest-vessel distance. At the individual vessel-segment level, we quantified six **structural parameters** and four **functional hemodynamics parameters**, hereafter referred to as **functional parameters**. The segment-level structural parameters were segment length, distance to the ONH periphery, depth relative to the anterior laminar insertion plane, tortuosity, inclination angle, and polar angle relative to the central retinal vessel center. The segment-level functional parameters were blood flow rate, blood pressure, blood oxygenation, and surrounding tissue oxygenation.

Finally, we developed a quadratic regression model to predict surrounding tissue oxygenation, which is difficult to measure experimentally, using experimentally measurable segment-level parameters. The predictors included the six segment-level structural parameters listed above together with blood flow rate. Below, we describe each step in more detail.

### 2.1 ONH vasculature reconstruction

#### Vessel labeling and histological preparation

We followed the process reported before.^38,43^ In accordance with University of Pittsburgh’s IACUC regulations and following NIH and ARVO guidelines for the use of animals, vessel labeling was performed on four healthy female rhesus macaque monkeys post-mortem within two hours of death. To regulate IOP during the procedure, the anterior chamber of each eye was cannulated, maintaining IOP at 5 mmHg. This pressure was selected to obtain the most complete vascular filling and labeling, without potential artifacts of an unphysiologic IOP of 0 or tissue distortions, including potential vessel constrictions, at higher IOPs. The goal was to label the entire vascular network rather than reproduce physiological hemodynamic conditions. Two polyimide microcatheters were inserted into the carotid arteries for vascular labeling. Warm phosphate-buffered saline (PBS) was initially perfused to flush the vascular bed, followed by the infusion of 100 ml of a lipophilic carbocyanine dye (Dil) into each carotid artery for vessel labeling. Fluorescence microscopy (FM) imaging confirmed strong signals in retinal and choroidal vessels, indicating successful labeling. Subsequently, 100 ml of 10% formalin was perfused to fix the tissues. The eyes were then enucleated and immersion-fixed in 10% formalin for 24 hours to ensure thorough preservation.

#### Tissue sectioning and imaging

The ONH and adjacent sclera were isolated using a 14-mm trephine. The tissues were cryoprotected and sectioned into 16 µm-thick slices for imaging. FM was used to visualize the labeled vessels, while instant polarized light microscopy (IPOL) was employed to visualize collagen structures.^44,45^ Sequential FM and IPOL images of every section were acquired, registered, and aligned using Avizo software (version 9.1; Thermo Fisher Scientific, MA, USA).

#### 3D vasculature reconstruction

The aligned FM images were segmented and skeletonized to generate a 3D reconstruction of the vasculature in the ONH. The reconstructed vascular network included the entire LC region, along with some pre-laminar and retro-laminar regions. The LC region was identified by the presence of collagen beams observed in the IPOL images.

Some non-uniform labeling, discontinuities, and leaks were sometimes observed. Manual corrections, including cleaning to remove artifact and bridging to close gaps, were applied as needed. We have discussed elsewhere^38,46^ the details of the labeling and reconstruction process and discussed the consequences and potential artifacts that may be introduced.

To assess reconstruction repeatability, two independent researchers reconstructed the same eye. Both were processed through the same computational pipeline to estimate hemodynamics and oxygenation. The reconstructions were highly consistent, with over 90% of vessel segments overlapping (Table 1). Differences in mean blood flow (2.96 nl/min vs. 2.61 nl/min) and mean oxygen partial pressure (55.56 mmHg vs. 54.85 mmHg) were small, indicating that manual corrections, including cleaning and bridging of discontinuous segments, had limited impact. These findings support the robustness and reproducibility of the reconstruction workflow, despite challenges associated with post-mortem labeling and manual intervention. A detailed comparison is provided in our previous work.^46^

**Table 1.** Repeatability of vascular reconstruction. Total and shared vessel segment counts, mean flow rate, and mean oxygen partial pressure are reported for two independent reconstructions of the same eye. The intraclass correlation coefficients between the reconstructions were 0.829 for flow rate and 0.985 for oxygen partial pressure.

|  | Total vessel segments | Shared vessel segments | Mean Flow rate (nl/min) | Mean Oxygen partial pressure (mmHg) |
|---|---|---|---|---|
| Reconstruction1 | 10918 | 9584 | 2.96 | 55.56 |
| Reconstruction2 | 10443 | 9584 | 2.61 | 54.85 |

#### Vessel diameter assumptions

Although vessel diameter data could potentially be extracted from the reconstructed images, post-mortem changes such as tissue swelling and pressure fluctuations may compromise the reliability of these measurements in reflecting in vivo conditions. Therefore, based on previous experimental data and consistent with other studies modeling ONH vasculature^29,47,48^, we assumed a uniform capillary diameter of 8 µm throughout the ONH region. Further studies are necessary to characterize vessel diameter under physiological conditions.

### 2.2 Structural and functional parameters definition

We defined structural parameters at two levels: five whole-ONH parameters describing the overall vascular geometry and six segment-level parameters describing individual vessel segments geometry.

The five whole-ONH structural parameters were total segment number, total branch-point number, ONH volume, total vessel length, and average nearest-vessel distance. A vessel segment was defined as the vascular path between two adjacent branch points, or between a branch point and a boundary endpoint. The average nearest-vessel distance was defined as the mean of the shortest distance from each ONH tissue point to the vascular network. This parameter provides a measure of the overall spatial coverage of the vascular network within the ONH.^49^

The six segment-level structural parameters were segment length, segment peripheral distance to the ONH canal wall, segment depth, tortuosity, inclination angle, and polar angle. These parameters were calculated for each vessel segment. An illustration of their definitions is shown in Figure 1, and the definitions were based on those introduced previously^38^.

<u>Segment peripheral distance</u> r was defined as the distance from the canal boundary to the midpoint of the vessel segment. <u>Segment depth</u> d was defined as the distance from the midpoint of the vessel segment to the reference plane defined by the anterior laminar insertion points. The <u>length</u> of each vessel segment l was measured as the pathway distance along the vessel from its start vertex to its end vertex.

#### Tortuosity

We defined tortuosity as the ratio of the path length of the vessel segment to the straight-line distance between its endpoints. A tortuosity value of one indicates that the vessel segment is straight, while larger tortuosity indicates a more twisted and curved segment.

#### Inclination angle

The inclination angle (0 ≤ θ ≤ π/2) of each vessel segment was defined as the angle between the end-to-end direction of each vessel and the anterior-posterior axis. An inclination of π/2 radians means that the vessel segment is aligned with the coronal plane.

#### Polar angle

Polar angle (0 ≤ φ ≤ π/2) was defined in the coronal plane using the central retinal vessels as the reference center. Specifically, it was calculated as the angle between the projected vessel-segment direction and the radial direction connecting the segment to the center of the central retinal vessels. When φ = 0, the vessel is aligned radially towards the canal center. When φ = π/2, the vessel is aligned circumferentially around the canal center.

These anatomical reference directions were selected to characterize vessel orientation relative to the peripheral-to-central blood supply and the laminar pattern of the LC, allowing us to investigate whether vascular alignment with these directions is associated with blood flow and tissue oxygenation. In addition, we performed a separate analysis of branch angles to characterize the relationships between local branching geometry and functional parameters. The branch angle was defined as the angle between two daughter branches at each branch point.

Four segment-level functional parameters were quantified: blood flow rate, blood pressure, blood oxygenation, and surrounding tissue oxygenation. <u>Blood flow rate</u> was defined for each vessel segment (nL/min). <u>Blood pressure</u> and <u>blood oxygenation</u> were averaged along each segment (mmHg). Tissue oxygenation is typically defined at tissue points. Here, to enable direct comparison with other segment-level parameters, we assigned a local tissue oxygenation value to each segment. <u>Surrounding tissue oxygenation</u> was defined as the average tissue oxygen partial pressure within 20 µm of the vessel surface (mmHg) (Figure 1C).

### 2.3 Hemodynamics and Oxygenation

For each of the four vessel networks we used the methodology described elsewhere to simulate the hemodynamics and calculate tissue oxygenation^39,41^. Briefly, blood pressure boundary conditions were applied to simulate the physiological blood supply for the ONH: Blood inflow from the circle of Zinn-Haller at the periphery, drainage through the central retinal vein, and interactions with the pre and retro lamina regions. Blood flow was modeled as incompressible laminar flow, described by the Poiseuille equation. Oxygen transport in the tissue was described by the diffusion-consumption equation.

#### Blood pressure boundary conditions

To simulate physiological blood supply to the ONH, pressure boundary conditions were assigned based on anatomical regions involved in blood supply and drainage: (Figure 1)

##### Periphery boundary

An arteriole pressure of 50 mmHg was set at the boundary vessel segment to represent blood inflow from the circle of Zinn–Haller.

##### Central boundary

The temporal half surface of the central vessel structure was defined as the central retinal vein boundary. A venous pressure of 15 mmHg was applied to the vessel segments on this surface to represent blood drainage through the central retinal vein.

The nasal half of the central vessel structure was defined as the central retinal artery boundary. A no-flux condition was imposed on vessel segments at this surface, indicating no blood exchange from the central retinal artery in or out.

##### Anterior boundary

The anterior surface interfaces with the prelaminar region, where tissue pressure was assumed equal to IOP (15 mmHg). A blood pressure of 20 mmHg was applied to vessel segments on this surface to ensure capillary patency under normal physiological conditions.

##### Posterior boundary

The posterior surface interfaces the retrolaminar tissue, where tissue pressure was estimated at 11 mmHg. A blood pressure of 16 mmHg was applied to the posterior boundary vessels to prevent collapse and allow sustained flow.

##### Blood flow modeling

We followed the approach described in^39^ to perform the ONH hemodynamics and oxygenation simulations. The ONH vascular network was modeled as a series of interconnected capillary segments joined at bifurcation points. Due to the low Reynolds number (< 0.05), blood flow within each capillary was approximated by Poiseuille’s flow^50^:

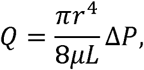

where Q is the flow rate, r is the vessel radius, L is the vessel length, *μ* is the blood viscosity, and Δ*P* is the pressure drop along the vessel.

Blood viscosity μ was described as a function of vessel diameter and hematocrit, incorporating the Fåhræus-Lindqvist effect. We used the empirical model proposed by Pries and Secomb^15^ to estimate effective viscosity. The detailed formulation and discussion can be found in^15^ and^51^. In our simulations, for a vessel diameter of 8 µm, the resulting effective viscosity was 7.65×10−3 Pa·s.

##### Oxygen transport in blood vessels

The oxygen flux in blood vessels satisfies:

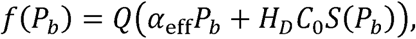

Where Q is the flow rate, *α*_eff_ is the effective solubility of oxygen in plasma, *H_D_* is the hematocrit, *C*_0_ is the concentration of hemoglobin-bound oxygen in a fully saturated red blood cell, *P_b_* is the blood oxygen partial pressure (mmHg), *S*(*P_b_*) is the oxygen-hemoglobin saturation, determined by the empirical oxygen–hemoglobin dissociation curve.

Oxygen conservation along each vessel segment gives,

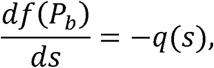

where *s* is arc-length parameter along the vessel, and *q*(*s*) is the oxygen flux per unit length through the vessel wall.

The diffusive oxygen flux across the cylindrical vessel wall is calculated from the radial gradient of tissue oxygen:

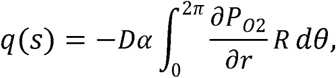

where *r* is the radial distance from the vessel centerline, R is the vessel radius, and the integral is around the circumference, denoted by angle *θ*.

##### Oxygen transport in neural tissue

Oxygen transport within the oxygen-consuming neural tissues was described using the reaction-diffusion equation^52^:

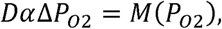

where D is the oxygen diffusion coefficient, α is the oxygen solubility coefficient, and *P_02_* is the tissue oxygen partial pressure. The oxygen consumption term *M*(*P_02_*) can be estimated by Michaelis-Menten enzyme kinetics^52^,

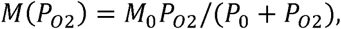

where *M*_0_ represents oxygen demand and *P*_0_ is the oxygen partial pressure at half-maximal consumption. In this study, oxygen demand was assumed uniform throughout ONH.

As reported before^39^, oxygen transport in the ONH was simulated using an efficient method combining an implicit finite-difference scheme with a multigrid solver for tissue oxygen, and ODEs for blood oxygenation along vessels. Vessel–tissue exchange was incorporated via a numerical delta function. All parameters used in the simulation are summarized in Table 2.

**Table 2.**
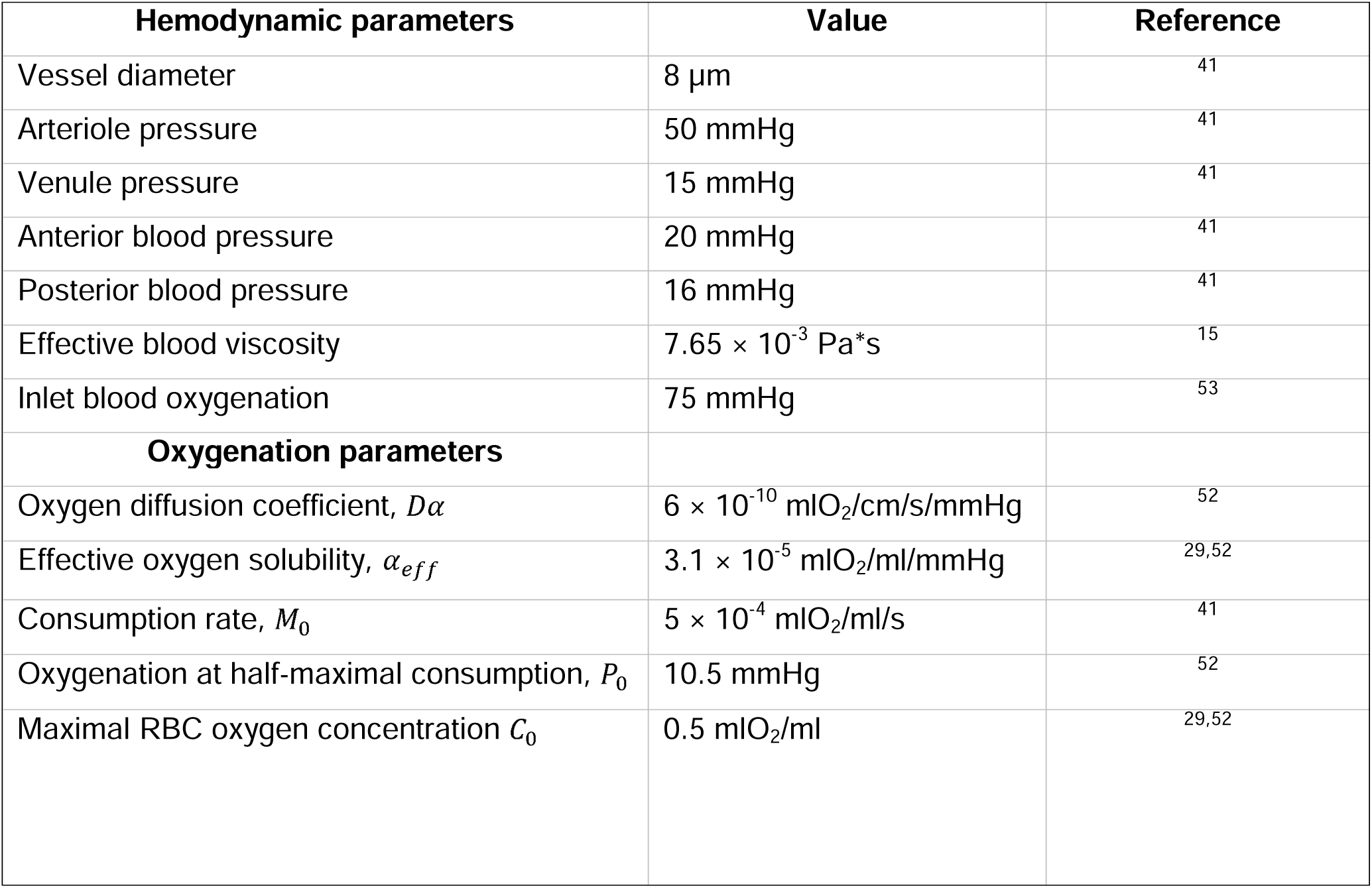
Parameters used in hemodynamic and oxygenation simulations.

### 2.4 Statistical analysis

Ten parameters were evaluated, including six structural parameters (peripheral distance, depth, length, tortuosity, inclination angle, and polar angle), and four functional parameters, including blood flow, blood pressure, oxygen saturation, and surrounding tissue oxygenation. Before statistical analysis all the parameters were normalized to have a mean of zero and a standard deviation of one. This allows for direct comparisons across parameters despite differences in units and/or scales.

Scatter plots were generated to visualize relationships between parameter pairs. For each pair, we considered linear, quadratic, logarithmic, and exponential models, selecting the curve with the highest R² value as the best fit. A Spearman correlation coefficient was calculated to quantify nonlinear associations, whereas a Pearson correlation coefficient was used for linear relationships. Additionally, histograms were created for each parameter to assess their distributions.

#### Multivariable mixed-effects model

We developed two multivariable mixed-effects models to estimate tissue oxygenation and hypoxia risk from experimentally measurable vascular features. Structural parameters and blood flow can be experimentally measured, whereas local tissue oxygenation remains difficult to measure directly. Therefore, the six structural parameters and blood flow were used as predictors in both models.

We employed a **multivariable quadratic mixed-effect regression model** to predict tissue oxygenation, which is crucial to axonal damage/disease progression. Although structural parameters and blood flow can be non-invasively measured in vivo for a limited number of vessels within the ONH, or comprehensively characterized ex vivo without flow information. Tissue oxygenation, however, remains difficult to measure directly. The objective of this model was to estimate tissue oxygenation from experimentally measurable vascular features. The model was applied to data from four eyes. The fixed-effect predictors included six structural parameter (depth (d_z_), periphery distance (d_p_), vessel segment length (l), tortuosity (*τ*), inclination angle (θ), and polar angle (φ) and one functional parameter, the logarithm of blood flow log(Q). The model was formulated as:

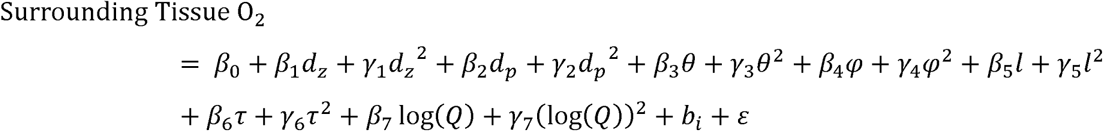

where β_k_ and *γ*_k_ are fixed effect coefficients for the linear and quadratic terms, respectively. 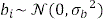 represents the random intercept for Eye i, and 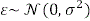 is the residual error term.

We then employed a **logistic mixed-effects model** to evaluate the probability of local hypoxia, defined as surrounding tissue O < 38 mmHg^42,54^, using the same vascular predictors as above. Unlike the linear formulation, which predicted continuous oxygenation, this model estimated the probability of hypoxia, P(O_₂_ < 38 mmHg), as a function of structural and flow parameters while accounting for eye-specific variability through a random intercept. The model was defined as:

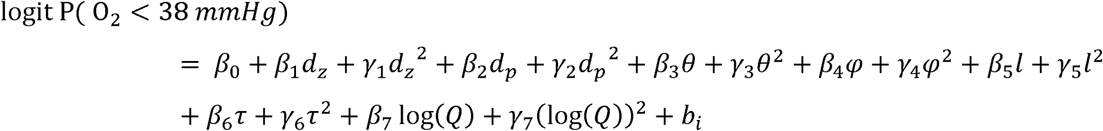

where logit(*ρ*) = ln [*ρ*/ (1-8)], β_k_ and *γ*_k_ are fixed-effect coefficients for the linear and quadratic terms, and 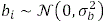 is the random intercept for Eye i. The model was fitted using maximum likelihood estimation under a binomial distribution with a logit link to quantify hypoxia risk from measurable vascular features.

To evaluate model performance, we used the Precision–Recall (PR) curve instead of the traditional ROC–AUC, since hypoxic regions (positive cases) were relatively rare in healthy tissue (< 10%). The PR curve better reflects performance on the positive class by illustrating how precision (the reliability of predicted hypoxia) varies with recall (the fraction of true hypoxic sites detected), providing a clearer measure of the model’s ability to identify hypoxic regions.

#### Spatial autocorrelation correction

Because measurements from nearby vessel segments may exhibit spatial autocorrelation, we performed a generalized least squares (GLS) sensitivity analysis following Dormann^55^. We incorporated a distance-dependent residual correlation structure to account for spatial dependence when estimating pairwise associations and their corresponding p values. Details of the procedure are provided below.

For each parameter pair, x_i_ and y_i_ denote the measurements for vessel segment i, and n denotes the number of segments. Each segment’s location was represented by its three-dimensional midpoint coordinates. The Euclidean distance between the segments i and j was denoted by d_ij_.

We constructed an *n* × *n* matrix C describing the correlations between regression residues.

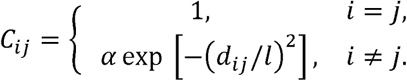

where α ɛ (0,1) represents the strength of spatial dependence and *l* represents its correlated distance scale. This specification allows stronger residual correlation between nearby segments, with correlation decreasing with distance. The parameters *α* and *l* were estimated by restricted maximum likelihood (REML), yielding the fitted correlation matrix C.

The residual for segment i was defined as

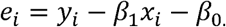

Using the distance correlation matrix C, GLS estimated these coefficients by minimizing the spatially weighted residual sum of squares:

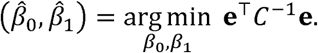

Where **e** = (e_1_, e_2_,…,e_n_)*^T^* is the residual vector, and 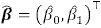 represents the estimated coefficients. The fitted GLS models provided estimates of pairwise associations and their corresponding p values. In the absence of spatial dependence *C* = 1, GLS reduces to ordinary linear least squares.

## 3. Results

Figure 2 illustrates the distribution of all parameters within a single ONH vascular network. The structural parameters, including polar angle, inclination angle, length, and tortuosity, showed generally weak correlations with depth and peripheral distance (|r| < 0.05, Figure 4), indicating a consistent vessel pattern throughout the network. Only inclination angle showed a weak positive association with depth (r ∼ 0.13). The hemodynamic and oxygenation parameters exhibited a radial-like spatial distribution, with higher flow rates, blood pressure, blood oxygenation, and tissue oxygenation in the peripheral region, and lower values in the central region. Table 3 shows the topological parameters across the eyes. The fitted distribution types and coefficients for all vessel structural and functional parameters across four eyes are provided in the supplementary material. A comparison of microvascular density and tortuosity between this work and other tissues and species is also shown in the supplementary material, evidence that our results are comparable to those from previous studies.

**Figure 2:**
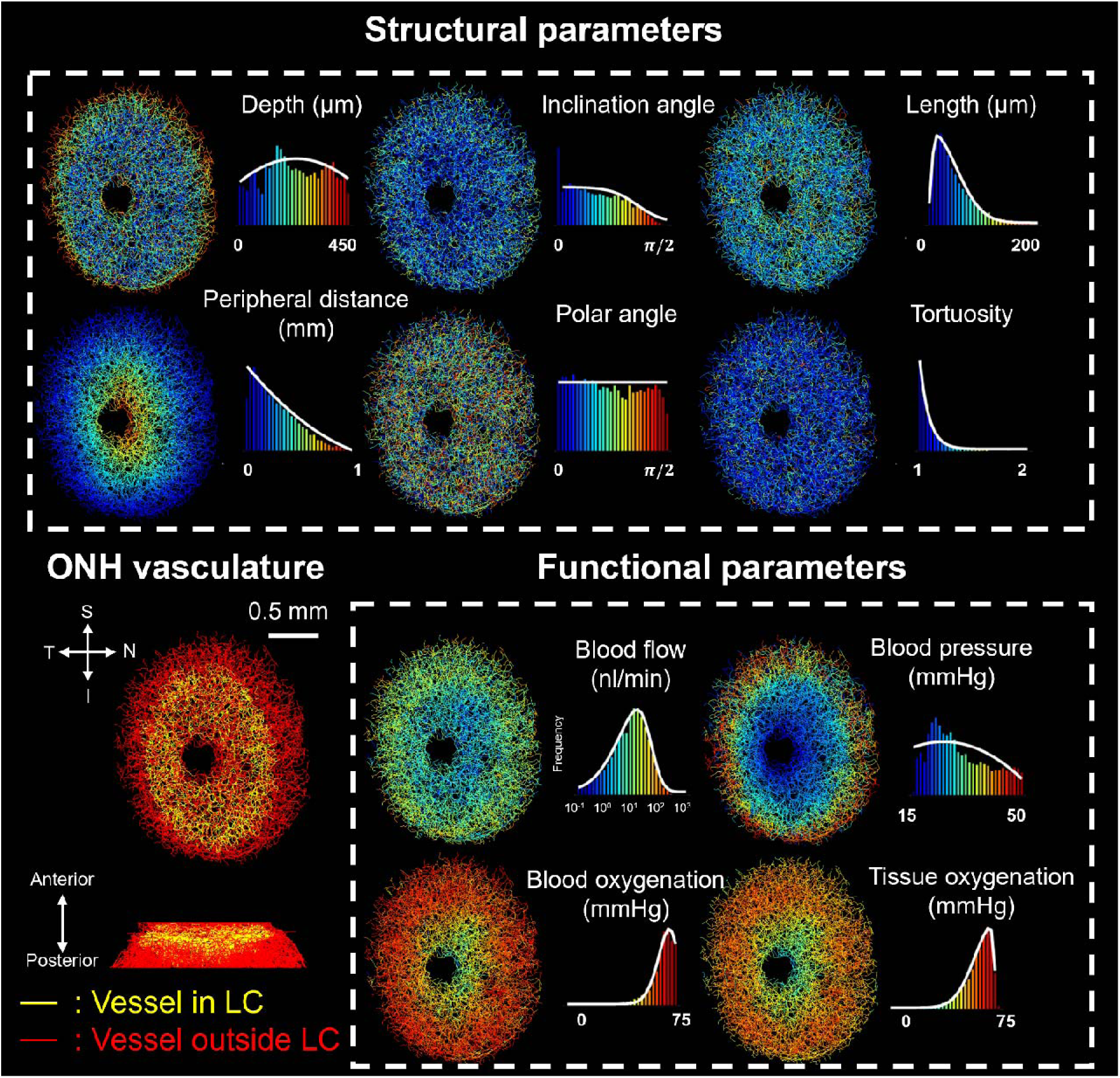
Distributions of structural and functional parameters in the ONH vasculature of a healthy monkey eye. The top panel illustrates structural parameters, including depth, peripheral distance, inclination angle, polar angle, segment length, and tortuosity, each represented with colored vasculature maps and corresponding histograms. The bottom panel provides functional parameters, showing blood flow, blood pressure, blood oxygenation, and tissue oxygenation with spatial distributions and histograms. Vascular orientation, length, and tortuosity, appear homogeneous across the ONH. Functional parameters displayed a radial spatial distribution. reflecting the ONH blood supply. Blood enters through the peripheral circle of Zinn-Haller and drains via the central retinal vein, creating pressure and oxygenation gradients from the periphery to the center. The ONH vasculature shown in the lower-left panel is from one of the eyes included in our previous work^42,46,54,59^, and is presented here using an adapted visualization to illustrate the overall geometry of the vasculature. Therefore, this panel may appear similar to the corresponding figure in our previous publication.

**Figure 3.**
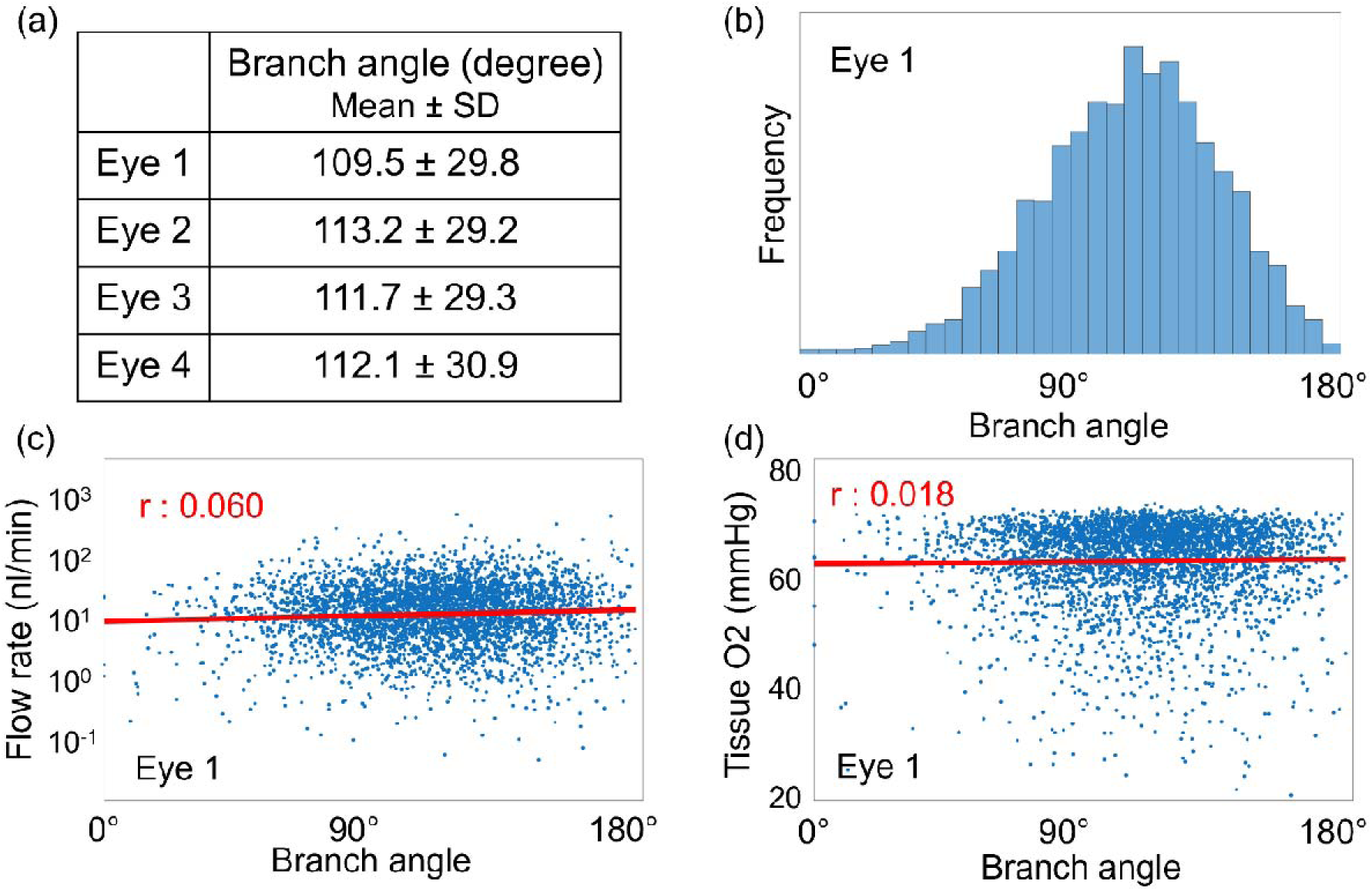
Branch angle analysis in the LC vascular network. (a) Mean ± standard deviation of branch angles across the four eyes were fairly similar. (b) Distribution of branch angles in Eye 1, showing a broad distribution with a peak around 110°–120°, reminiscent of the 120° configuration of three evenly spaced branches in a plane. (c–d) Relationships between branch angle and parent-vessel flow rate (c) and surrounding tissue oxygen tension (d) in Eye 1. Each dot represents a branch point, with the corresponding parent-vessel flow rate and local tissue oxygen tension. Red lines indicate linear regression fits. Pearson correlation coefficients were r = 0.060 (p = 0.00143) for flow rate and r = 0.018 (p = 0.242) for tissue oxygen tension, indicating negligible linear associations with branch angle.

**Figure 4:**
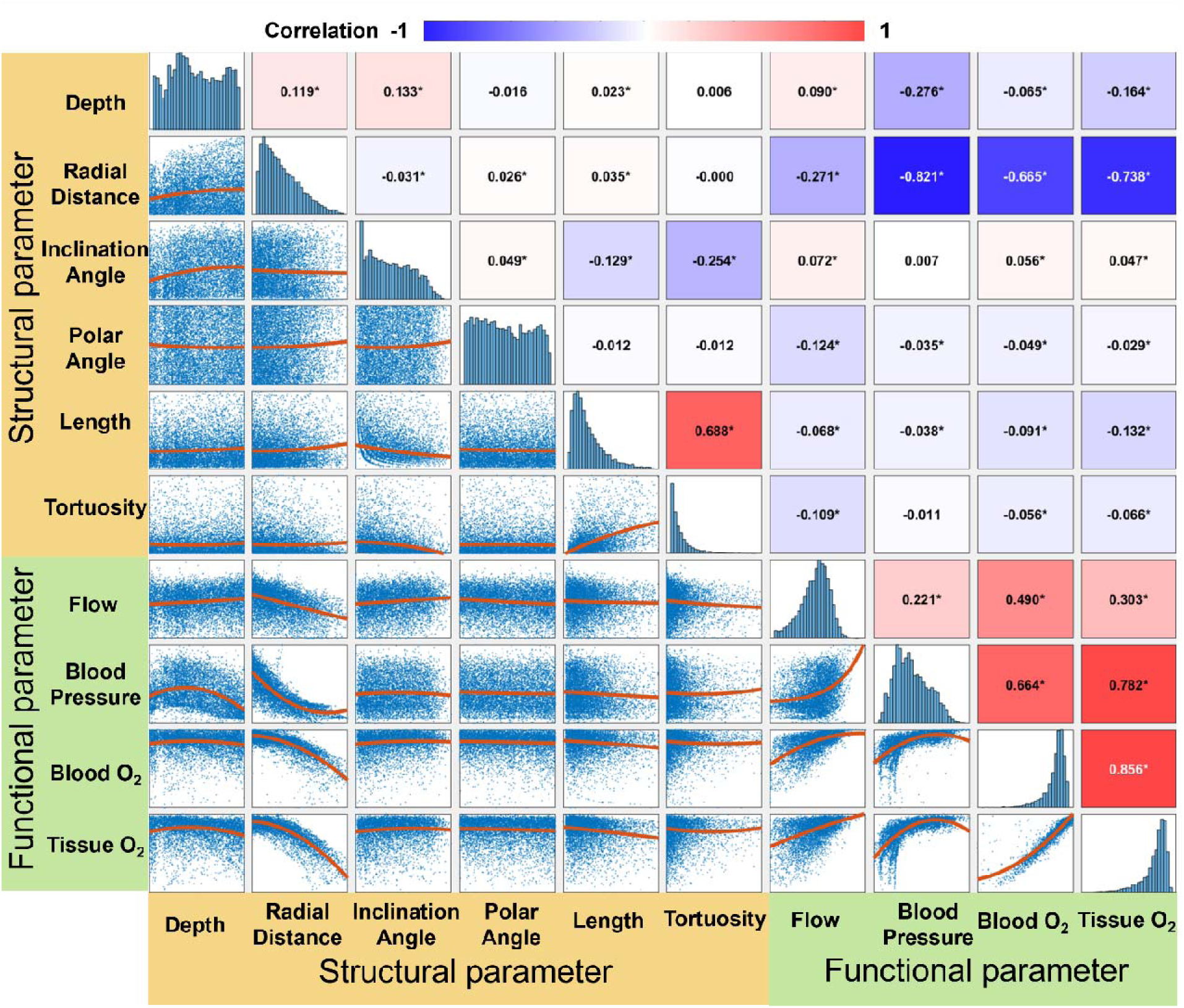
Scatterplots, distributions and correlations between parameters for Eye 1. Each dot in the scatterplots corresponds to a vessel segment. The red line in each scatterplot represents the best-fitting function. To help visualize trends, the correlation values are colored using blue to indicate strong negative correlations and red for strong positive correlations, with white indicating no correlation. Asterisks (*) indicate statistical significance (*p* < 0.05). Most correlations (38 of 45) were significant. Several weak associations involving polar angle, vessel length, or tortuosity had *p* ≥ 0.05. The strongest positive correlation (0.856) was observed between the blood oxygenation and tissue oxygenation. Flow and tissue oxygenation showed a moderate positive correlation (0.303), indicating the alignment between the two parameters, although potential mismatches or divergences might occur. Correlations between vessel orientation (inclination and polar angles) and functional parameters were minimal (< 0.1), suggesting that radial and circumferential vessels exhibit similar flow and oxygenation patterns, despite the overall radial distribution of these functional parameters. Depth and peripheral distance, as indicators of spatial location, exhibited relatively stronger correlations, suggesting that positional parameters may have a greater impact on functional behavior than local geometric features.

**Table 3.** Parameters for ONH vasculature topology across all eyes, including on the total segment number, vertex number, ONH volume (mm³), total vessel length (mm), and the average nearest vessel distance (µm). The segment number exhibits the greatest variation, while the average nearest vessel distance shows the least variation. This suggests that, despite differences in network size and volume, the average nearest vessel distance, which reflects tissue irrigation by blood vessels, remains relatively consistent across all eyes.

|  | Segment number | Branch point number | Volume (mm<sup>3</sup>) | Total vessel length (mm) | Average nearest vessel distance (μm) |
|---|---|---|---|---|---|
| Eye1 | 12966 | 9089 | 1.456 | 646.6 | 53.47 |
| Eye2 | 10918 | 8886 | 1.304 | 528.7 | 52.54 |
| Eye3 | 12012 | 9436 | 1.298 | 547.8 | 56.03 |
| Eye4 | 10700 | 8151 | 1.169 | 628.3 | 61.62 |

We further investigated the relationship between branch angle and vascular function (Figure 3). The average branch angles were around 110° across the four eyes. In the hierarchical retinal vasculature, larger parent vessels divide into smaller daughter vessels, and the reported mean angle between the two daughters is approximately 80°^56–58^. Vessel segments at LC branch points tend to have similar radii, which may help explain why the distribution peaks near 120°, the angle between three evenly spaced branches in a plane. In Eye 1, branch angle showed negligible linear correlations with parent-vessel flow rate (r = 0.060) and surrounding tissue oxygen tension (r = 0.018) (Figure 3c–d). Across the four eyes, these correlations were negligible in magnitude, with only one being statistically significant.

Scatterplots, distributions, and pairwise correlations between structural and functional parameters are summarized in Figure 4. Functional parameters, particularly blood and tissue oxygenation, showed strong correlations, while structural-functional associations were generally weak. Spatial location parameters such as depth and peripheral distance showed stronger correlations with tissue oxygenation than vessel-specific geometric features such as orientation, suggesting that vessel function is more influenced by its position within the ONH than by its own geometry.

Figure 5 summarizes correlations between vessel parameters and tissue oxygenation across four eyes. While correlation directions were generally consistent, morphological parameters (e.g., length, tortuosity, orientation) exhibited weak and variable correlations with tissue oxygenation. In contrast, spatial and functional parameters showed stronger and more consistent correlations, suggesting their importance for vessel-level oxygenation prediction.

**Figure 5.**
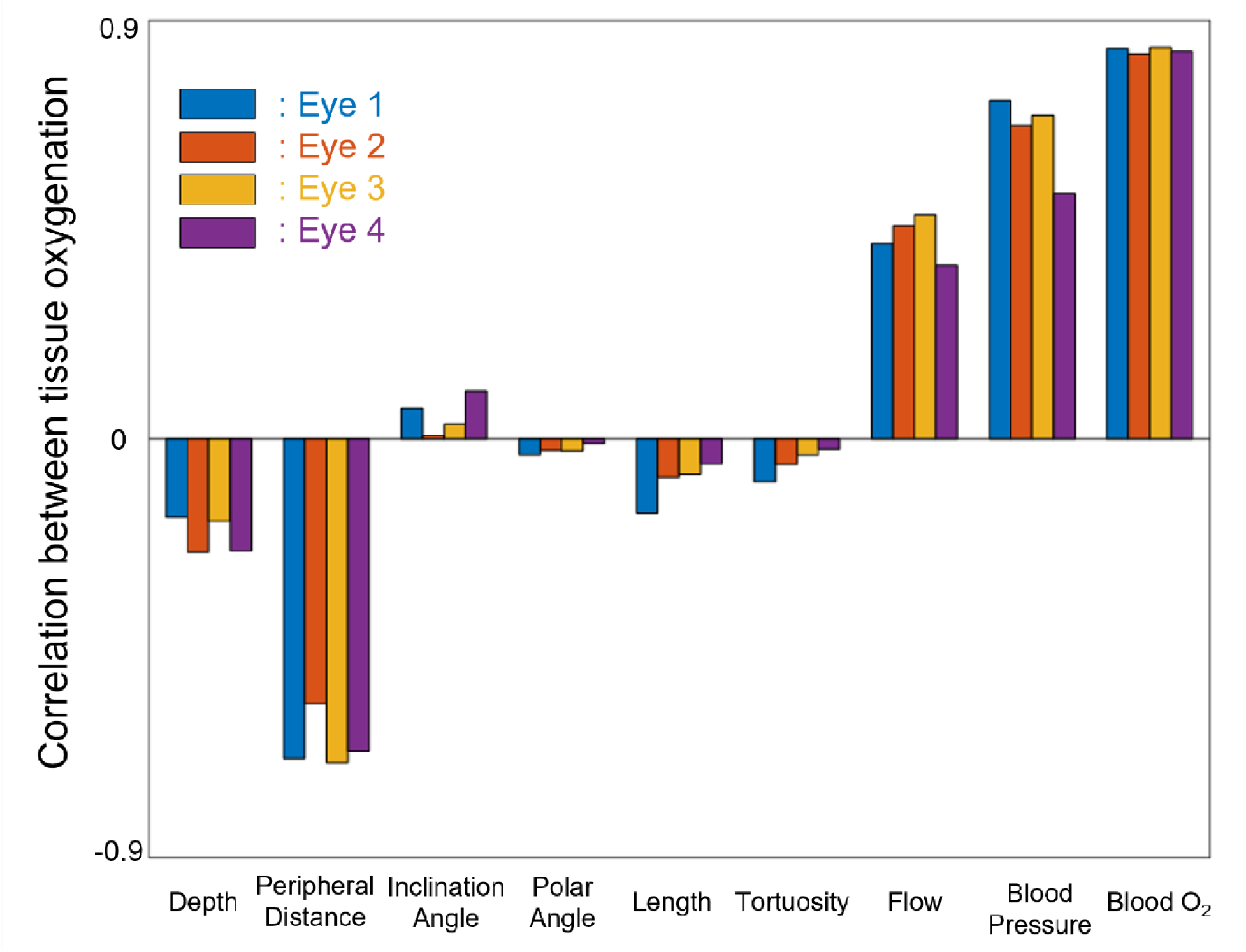
Correlations between all the parameters and tissue oxygenation across four eyes. Each parameter exhibited a consistent positive or negative correlation with tissue oxygenation across all eyes, with generally similar correlation strengths. However, the strength of these correlations varied substantially among parameters, particularly for vessel morphological parameters such as length, tortuosity, and orientation, which generally showed weak associations. This variability suggests that morphological features alone may be insufficient to accurately predict tissue oxygenation at the single-vessel level. In contrast, spatial and functional parameters, including depth, peripheral distance, flow, and pressure, showed stronger and more consistent correlations, suggesting that such parameters may be essential for the reliable prediction of tissue oxygenation at the vessel level.

Although blood flow is commonly used as an indicator of hemodynamic function and is significantly correlated with tissue oxygenation (r ≈ 0.313, p < 0.001), Figure 6 reveals limitations of relying on flow alone for the ONH. Several segments show substantial differences from the expected flow–oxygenation relationship, particularly within the central ONH. These findings indicate that blood flow alone does not provide a reliable assessment of tissue oxygenation and support the need for more comprehensive models that incorporate additional structural and spatial parameters to improve the prediction of oxygenation.

**Figure 6.**
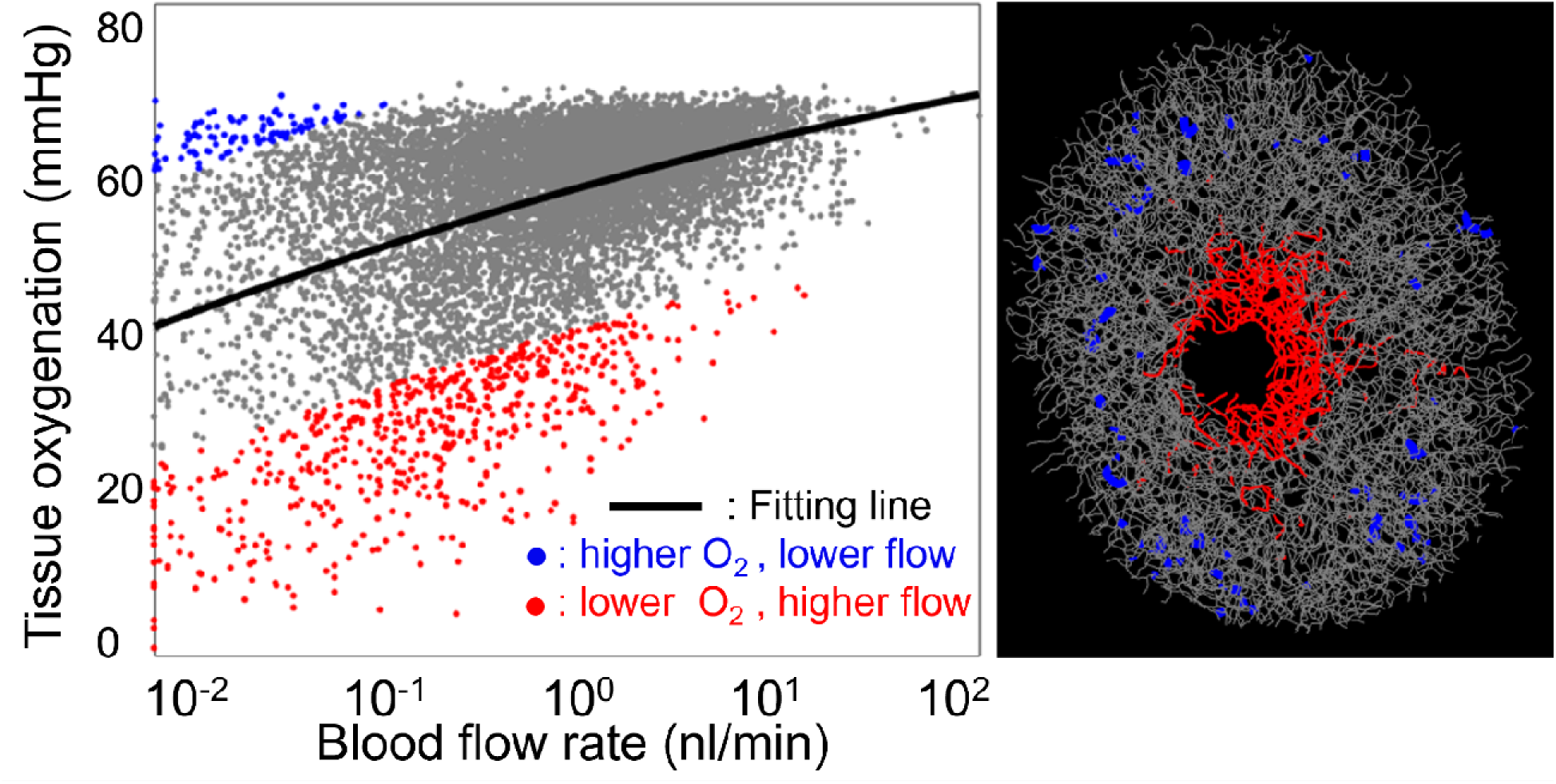
Mismatch region between blood flow rate and tissue oxygenation in Eye 1. **(Left)** Scatterplot of blood flow rate versus tissue oxygenation. Each point corresponds to a vessel segment. Vessels with higher-than-expected oxygenation relative to flow (blue) were defined as having oxygenation values ≥15 mmHg above the fitting line, while those with lower-than-expected oxygenation relative to flow (red) were ≥15 mmHg below it. Most vessel segments (gray) were located near the fitting line, indicating a general consistent flow-oxygenation relationship. **(Right)** Spatial distribution of mismatch segments. Vessels with lower-than-expected oxygenation relative to flow w were concentrated in the central ONH, likely reflecting its downstream position in the vascular network, where oxygen is depleted despite sustained flow. These mismatches were more common on the temporal (CRA) side, which lacks flow exchange with adjacent ONH regions. Higher-than-expected oxygenation relative to flow was less common and distributed around the ONH.

Table 4 summarizes the fitted coefficients from the multivariable quadratic mixed-effects model used to predict tissue oxygenation. The model demonstrated strong performance (R² = 0.75) and identified peripheral distance, blood flow, and depth as the most influential predictors. These results emphasizing the combined role of functional and spatial factors in determining ONH tissue oxygenation.

**Table 4.** Summary of coefficients from the multivariable quadratic mixed-effects model relating vessel parameters to predicted tissue oxygenation. Both original and normalized estimates are provided to allow for physiological interpretation and comparison of relative importance. The model includes linear and quadratic terms for seven predictors. Original estimates are expressed in physical units, while normalized estimates correspond to standardized variables (mean = 0, Std = 1). Among all predictors, peripheral distance, depth, and blood flow were the most influential in determining tissue oxygenation.

|  | Original Estimate | Normalized Estimate | pValue |
|---|---|---|---|
| Intercept | 45.3 ± 1.55 | 0.437 ± 6.70*10<sup>-3</sup> | <10<sup>-5</sup> |
| **Linear Term Coefficient** |  |  |  |
| Depth (β<sub>1</sub>, mmHg/μm) | 0.129 ± 1.56*10<sup>-3</sup> | -0.0758 ± 2.99*10<sup>-3</sup> | <10<sup>-5</sup> |
| Peripheral distance (β<sub>2</sub>, mmHg/μm) | -0.0116 ± 5.74*10<sup>-4</sup> | -0.606 ± 3.80*10<sup>-3</sup> | <10<sup>-5</sup> |
| Inclination angle (β<sub>3</sub>, mmHg/rad) | 0.735 ± 0.244 | 0.0174 ± 3.21*10<sup>-3</sup> | 2.6*10<sup>-3</sup> |
| Polar angle (β<sub>4</sub>, mmHg/rad) | 0.347 ± 0.244 | 1.01*10<sup>-3</sup> ± 2.83*10<sup>-3</sup> | 0.16 |
| Length (β<sub>5</sub>, mmHg/μm) | -0.0251 ± 2.64*10<sup>-3</sup> | -0.0761 ± 4.42*10<sup>-3</sup> | <10<sup>-5</sup> |
| Tortuosity (β<sub>6</sub>, mmHg) | 9.28 ± 1.66 | 0.0354 ± 4.85*10<sup>-3</sup> | <10<sup>-5</sup> |
| log(Flow) (β<sub>7</sub>, mmHg/log(nl/min)) | 1.61 ± 0.0283 | 0.182 ± 3.43*10<sup>-3</sup> | <10<sup>-5</sup> |
| **Quadratic Term Coefficient** |  |  |  |
| Depth (γ<sub>1</sub>, mmHg/μm<sup>2</sup>) | -2.94*10<sup>-4</sup> ± 3.35*10<sup>-6</sup> | -0.271 ± 2.90*10<sup>-3</sup> | <10<sup>-5</sup> |
| Peripheral distance (γ<sub>2</sub>, mmHg/μm<sup>2</sup>) | -3.41*10<sup>-5</sup> ± 7.28*10<sup>-7</sup> | -0.131 ± 2.54*10<sup>-3</sup> | <10<sup>-5</sup> |
| Inclination angle (γ<sub>3</sub>, mmHg/rad<sup>2</sup>) | -0.286 ± 0.195 | -6.20*10<sup>-3</sup> ± 2.78*10<sup>-3</sup> | 0.14 |
| Polar angle (γ<sub>4</sub>, mmHg/rad<sup>2</sup>) | -0.196 ± 0.152 | -4.13*10<sup>-3</sup> ± 3.16*10<sup>-3</sup> | 0.20 |
| Length (γ<sub>5</sub>, mmHg/μm<sup>2</sup>) | 2.84*10<sup>-5</sup> ± 1.60*10<sup>-5</sup> | -2.97*10<sup>-4</sup> ± 1.91*10<sup>-3</sup> | 0.08 |
| Tortuosity (γ<sub>6</sub>, mmHg) | -2.99 ± 0.618 | -4.19*10<sup>-3</sup> ± 1.53*10<sup>-3</sup> | <10<sup>-5</sup> |
| log(Flow) (γ<sub>7</sub>, mmHg/log<sup>2</sup>(nl/min)) | -0.189 ± 0.0167 | -0.0212 ± 2.20*10<sup>-3</sup> | <10<sup>-5</sup> |
| **Random effect** |  |  |  |
| Random intercept SD (σ<sub>b</sub>) | 2.25 |  |  |
| Residual SD (σ) | 5.04 |  |  |

**R<sup>2</sup> = 0.75**

Table 5 and Figure 7 summarize the results of the logistic mixed-effects model for hypoxia classification. The fitted coefficients highlight the contributions of structural and flow parameters to hypoxia risk. The precision–recall AUC of 0.807 indicates good classification performance, demonstrating that the model can distinguish regions at risk of hypoxia from well-oxygenated tissue using localized vascular information.

**Figure 7.**
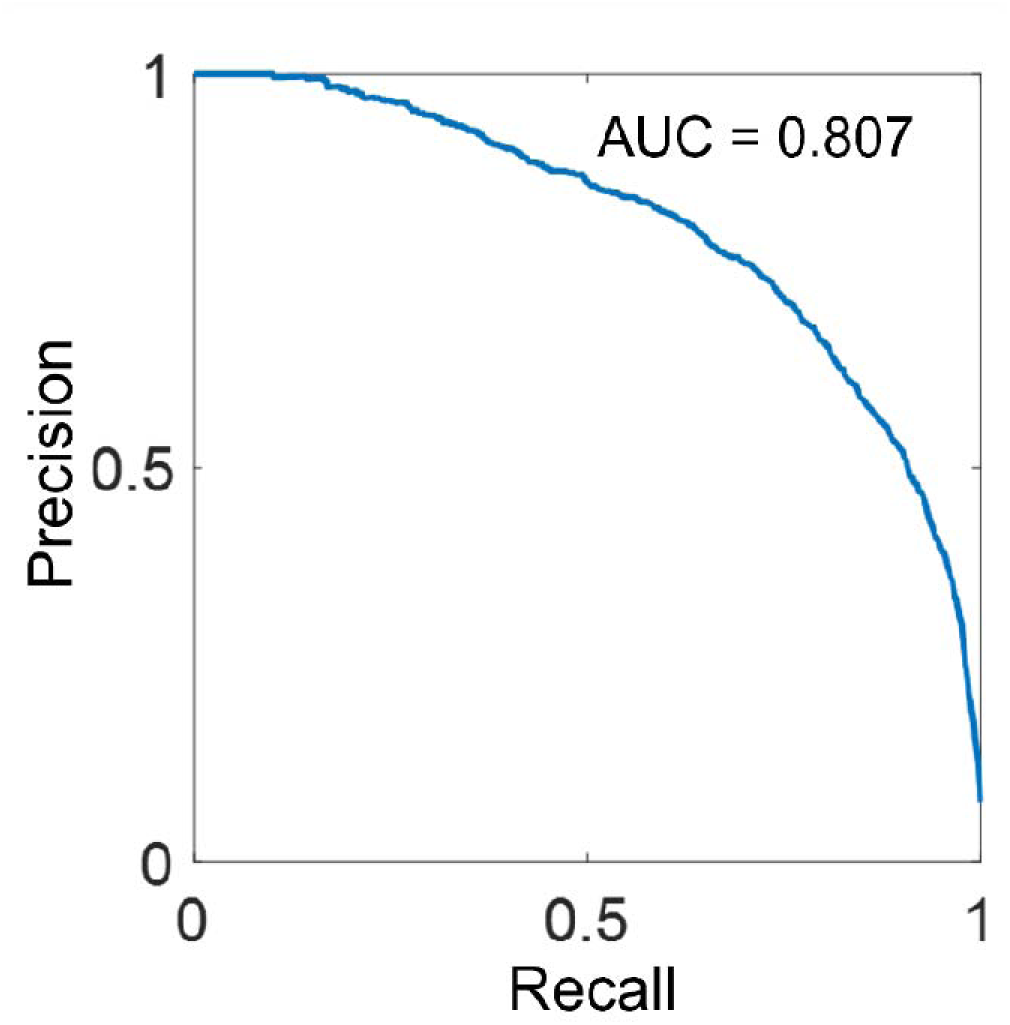
Precision–recall curve for the logistic mixed-effects model. The positive case represents hypoxia (O₂ < 38 mmHg). Precision is defined as true positive/(true positive + false positive), and recall as true positive/(true positive +false negative). With a precision– recall AUC of 0.807, the model showed good sensitivity and reliability in identifying hypoxic regions.

**Table 5.** Summary of coefficients from the quadratic logistic mixed-effects model relating vessel parameters to the probability of hypoxia. Both original and normalized estimates are provided to show physiological interpretation and comparison of relative importance. Among all predictors, peripheral distance, depth, and blood flow were the most influential factors in predicting hypoxia probability.

|  | Original Estimate | Normalized Estimate | pValue |
|---|---|---|---|
| Intercept | -7.36 ± 0.450 | -7.65 ± 0.588 | <10<sup>-5</sup> |
| **Linear Term Coefficient** |  |  |  |
| Depth (β<sub>1</sub>, mmHg/μm) | -0.420 ± 0.0339 | -0.290 ± 0.0335 | <10<sup>-5</sup> |
| Peripheral distance (β<sub>2</sub>, mmHg/μm) | 2.27± 0.0991 | 2.48 ± 0.103 | <10<sup>-5</sup> |
| Inclination angle (β<sub>3</sub>, mmHg/rad) | -0.127 ± 0.0382 | -0.156 ± 0.0380 | <10<sup>-5</sup> |
| Polar angle (β<sub>4</sub>, mmHg/rad) | 0.0450 ± 0.0325 | 0.0577 ± 0.0330 | 0.08 |
| Length (β<sub>5</sub>, mmHg/μm) | 0.0806 ± 0.0501 | 0.0669 ± 0.0489 | 0.17 |
| Tortuosity (β<sub>6</sub>, mmHg) | -0.0868 ± 0.0588 | -0.0982 ± 0.0523 | 0.06 |
| log(Flow) (β<sub>7</sub>, mmHg/log(nl/min)) | -0.762 ± 0.0801 | -0.704 ± 0.0735 | <10<sup>-5</sup> |
| **Quadratic Term Coefficient** |  |  |  |
| Depth (γ<sub>1</sub>, mmHg/μm<sup>2</sup>) | 1.24 ± 0.0357 | 1.28 ± 0.0361 | <10<sup>-5</sup> |
| Peripheral distance (γ<sub>2</sub>, mmHg/μm<sup>2</sup>) | 0.330 ± 0.0395 | 0.274 ± 0.0394 | <10<sup>-5</sup> |
| Inclination angle (γ<sub>3</sub>, mmHg/rad<sup>2</sup>) | 0.0497 ± 0.0339 | 0.0583 ± 0.0325 | 0.07 |
| Polar angle (γ<sub>4</sub>, mmHg/rad<sup>2</sup>) | -0.0336 ± 0.0365 | -0.0481 ± 0.0368 | 0.19 |
| Length (γ<sub>5</sub>, mmHg/μm<sup>2</sup>) | 0.0239 ± 0.0207 | 0.0325 ± 0.0202 | 0.11 |
| Tortuosity (γ<sub>6</sub>, mmHg) | 0.0178 ± 0.0213 | 0.0205 ± 0.0169 | 0.22 |
| log(Flow) (γ<sub>7</sub>, mmHg/log<sup>2</sup>(nl/min)) | -7.13*10<sup>-4</sup> ± 0.0363 | 0.0387 ± 0.0340 | 0.26 |
| **Random effect** |  |  |  |
| Random intercept SD (σ<sub>b</sub>) | 1.14 |  |  |

## 4. Discussion

We leveraged 3D eye-specific numerical models to characterize the structure, blood supply and oxygenation of the ONH vascular network. Specifically, we investigated the distributions and correlations among the structural, hemodynamics, and oxygenation parameters. Four main findings arise from the work: (1) The structural parameters did not show any clear spatial patterns, while the hemodynamics/oxygenation parameters exhibited a radial-like spatial distribution. (2) Blood flow, blood pressure, blood oxygenation and tissue oxygenation generally exhibited positive correlations, but diverge at certain regions. (3) Vessel length and tortuosity showed weak negative correlations with functional parameters, while vessel orientation exhibited negligible associations. (4) Multivariable mixed-effects modeling identified blood flow and spatial parameters as strong predictors of tissue oxygenation.. Detailed discussions on these findings are provided below.

### The structural parameters did not show any clear spatial patterns, while the hemodynamics/oxygenation parameters exhibited a radial-like spatial distribution

Vessel length, tortuosity, polar angle, and inclination angle generally showed weak correlations with depth and peripheral distance (|r| < 0.05), with only inclination angle showing a weak positive association with depth (r ∼ 0.2). This weak correlation indicates that vessels are highly random and evenly distributed throughout the ONH vasculature. Consequently, the vascular network appears homogeneous across different regions of the ONH, implying that each location within the ONH is supplied by a similar vascular pattern. Such geometric uniformity of ONH vasculature could contribute to a smooth and widespread irrigation of blood and oxygen across the entire ONH, which crucial for the health of retinal ganglion axons within.

Functional parameters displayed a radial spatial distribution. The radial patterns can be explained from the structure of the ONH blood supply, Blood enters the ONH through the circle of Zinn-Haller, a peripheral vascular ring, and flows toward the central retinal vein for drainage.^13^ This inflow-outflow dynamic creates a gradient of pressure and oxygenation from the periphery towards the central ONH.

### Blood flow, blood pressure, blood oxygenation and tissue oxygenation generally exhibit positive correlations, but diverge at certain regions

All the functional parameters, blood flow, blood pressure, blood oxygen and tissue oxygen show positive correlations. Vessels with higher flow rates generally tend to have higher blood pressure, higher blood oxygenation saturation, and higher surrounding tissue oxygenation. Higher blood pressure typically indicates an upstream vessel, such as an arteriole, and is associated with higher flow and oxygenation, as observed in other organs.^60–62^ However, in the case of ONH, flow rate and oxygenation can diverge in certain regions. Specifically, the central region of ONH, which is considered a downstream or outflow region, has the lowest oxygenation but high flow rate.

Blood perfusion rate has long been considered a valuable clinical biomarker^13,63–65^, largely due to limited direct oxygenation measures. However, our results highlight the importance of distinguishing blood flow and oxygenation. Although the blood flow has its own physiological importance, including removal of metabolic waste, hormone regulation, immune response, and fluid balance, the fundamental biological function of the circulation system is to deliver oxygen, with blood flow serving as the transport medium. Tissue oxygenation is therefore a more direct and essential measure of cellular activity and metabolic support, particularly in the ONH, which contains dense neural tissues with high oxygen demands. Oxygenation appears to be the most relevant measure to predict or understand glaucomatous neuropathy. The different behaviors of blood perfusion and oxygenation in this study suggests that blood flow may not be an adequate measure, and could potentially under/overestimate the oxygen supply in certain ONH regions.

### Vessel length and tortuosity showed weak negative correlations with functional parameters, while vessel orientation exhibited negligible associations

Our results showed that vessel length and tortuosity have weak negative correlations with both blood flow and oxygenation (∼-0.1). Long and tortuous vessels tend to have lower blood flow and oxygenation. This can be explained by flow resistance in each vessel segment, which is influenced by vessel length and tortuosity. Long and twisted vessels exhibit higher flow resistance^50^, limiting blood transport efficiency and reducing oxygenation.

The frequency distribution of vessel characteristics shows that most vessels in the ONH are short and straight, following an exponential-like distribution, with approximately 90% of vessels shorter than 5 µm and with tortuosity less than 1.2. This could result from the high demand from the neural tissues in ONH region. A dense vascular network with short and straight vessels could support blood and oxygen to ONH region efficiently.

It is important to emphasize that the negative relationship is weak. Although vessel length and tortuosity influence flow conductance for single segment, the highly interconnected ONH vasculature makes flow and oxygenation interdependent across the network. Blood flow and oxygenation in each vessel segment result from cumulative effects from upstream segments and the overall network, including loops, bifurcations, and parallel branches. Therefore, if a small, localized area contained only long and tortuous vessels, it could still be irrigated by surrounding regions. However, if the ONH were to experience widespread loss of short capillaries, perfusion and oxygen supply would likely be compromised, as discussed in^42^.

Beyond the segment geometry, one may also expect that vessel orientation to influence functional parameters. Radial vessels could have higher flow rates and oxygenation due to the radial perfusion pattern in ONH. However, our findings revealed only weak associations between these parameters. This could be explained from the specific biological function of the ONH vasculature: providing adequate blood transport through and providing sufficient oxygenation to the neural tissue within. This dual function is reflected in the topology and structure of ONH vasculature.

Efficient transport typically relies on tree-like, hierarchical vessel networks, that concentrate materials on main branches, reducing redundancy and optimizing transport efficiency.^17,18^ Such structures are observed in many vascular systems focused on transport, like the kidney arterial tree^66^. In contrast, irrigation efficiency depends on dense, interconnected microvascular networks that enable extensive exchange with surrounding tissues. These networks include “redundant” vessels, which enhance robustness by providing alternative pathways for blood flow and improving oxygen by bringing vessels closer to high-demand tissues, as observed in capillary systems focused on tissue exchange.^67^

In the ONH, radial direction vessels are essential for transport due to the radial perfusion pattern. However, similar flow rates in both radial and circumferential vessels suggest that the ONH vasculature primarily focuses on neural tissue irrigation rather than pure transport efficiency. This structure could provide robustness against vascular challenges, such as stochastic and IOP-related vascular deformation, as discussed in our previous work.^42^

### Multivariable mixed-effects modeling identified blood flow and spatial parameters as key predictors of tissue oxygenation

Our multivariable mixed model achieved strong performance (R² = 0.75) in predicting tissue oxygenation based on experimentally measurable parameters. Peripheral distance, blood flow, and depth emerged as the most influential predictors, reflecting the key roles of spatial positioning and local hemodynamics in determining tissue oxygenation. The strong dependence on spatial parameters, particularly peripheral distance, can be attributed to the blood flow pattern in the ONH, where peripheral vessels are typically upstream and thus associated with higher flow and oxygenation levels. In contrast, vessel orientation had a relatively high p-value, indicating limited contribution, consistent with its weak correlation with functional parameters. Given the challenges associated with direct measurement of tissue oxygenation, estimation based on measurable structural, spatial, and flow parameters may provide a practical approach for assessing local oxygenation. Although the model does not account for cumulative flow and oxygen transport across the entire vascular network, incorporating vascular connectivity and upstream transport dynamics may further improve predictive accuracy. Overall, these findings support the feasibility of estimating local ONH oxygenation from limited measurable vascular information.

To our knowledge, this is the first study to integrate structural and functional analyses of eye-specific ONH vasculature, focusing on vascular morphology, hemodynamics, and oxygenation. Previous analyses primarily addressed overall ONH morphology without detailing vascular structure and function. Our method reconstructs high-resolution ONH vascular networks from histological sections of healthy monkey eyes, ensuring detailed vascular geometry information. By combining these reconstructions with accurate and efficient numerical simulations, we are able to predict the ONH hemodynamics and oxygenation at the capillary segment, enabling a systematic analysis of key structural and functional parameters across multiple eyes.

### Limitations

It is important to acknowledge the limitations of this study so that readers can consider them when interpreting our findings. One major one is that we set the diameter of the vessel as 8 µm in this work. The rationale and limitations are discussed in detail in the method section. We want to highlight that vessel diameter has been identified as a critical parameter influencing hemodynamics and oxygenation in the ONH in previous studies.^41,68^ We expect a positive correlation between vessel segment diameter and functional parameters, which we will examine further once more reliable in vivo vessel diameter data becomes available.

An important limitation of our study is the lack of blood flow autoregulation in the ONH model. Our model assumes that all parameters are time-invariant, whereas in vivo, blood flow and oxygen supply are dynamically regulated in response to changes in physiological conditions and metabolic demands.^69–71^ Through autoregulation, blood vessels can constrict or dilate in response to changes in perfusion pressure, metabolic demand, and local signals such as hypoxia. These responses adjust vascular resistance and blood flow distribution to help maintain adequate tissue perfusion and oxygenation.^69^ Although the exact mechanisms of autoregulation and vascular remodeling in the ONH remain unclear, these processes influence blood flow and oxygen supply,^64,69,71–74^ and may affect the associations between vascular structure and function as identified in this study. Our results should therefore be interpreted as model-based predictions under fixed physiological conditions rather than a complete representation of in vivo hemodynamics. Future studies incorporating dynamic autoregulatory mechanisms are necessary for more complete understanding of ONH structure–function relationships and their potential roles in diseases such as glaucoma.

Another limitation is that we consider the functional parameters of each vessel segment independently in our statistical analysis, despite the interdependent flow and oxygenation across the highly connected ONH vascular network. Hemodynamics in one segment is influenced by the entire ONH vasculature; for example, surrounding vessels contribute to a segment’s perfusion and oxygenation, and longer-distance effects, such as reduced downstream perfusion due to upstream narrowing vessels, also play a significant role. A consequence of this assumption is that the statistical significance of the parameter associations will be inflated. Future study can account for correlations within a parameter (usually called autocorrelations) to remove this effect and improve the statistical analysis^75^.

Also importantly, although another primary function of ONH is to provide mechanical support to the axons and vessels within, we did not incorporate the biomechanical characteristics, such as pressure-induced vessel deformations into our analysis. Our model was developed using healthy eyes with normal perfusion and oxygenation, and does not reflect pathological scenarios such as compromised perfusion under elevated IOP. While we included detailed eye-specific vascular networks, we did not consider the corresponding collagenous structure of LC beams and pores. Recent studies show that vascular and connective tissue networks are distinct, suggesting a more complex structure-function relationship.^12,43^ Future work will incorporate both biomechanical and vascular components under pathological conditions to better understand the role of structural parameters in development of glaucoma.

Finally, the oxygen transport model relies on several simplifying assumptions regarding vascular hemodynamics, tissue oxygen transport, and metabolic consumption. The physiological properties of the LC, including metabolic activity, tissue oxygen diffusivity, and oxygen exchange between blood vessels and surrounding tissue, are not yet fully characterized. These parameters may influence the predicted oxygen distribution and the quantitative accuracy of the model. Experimental validation is needed to assess the accuracy and reliability of the LC oxygen transport model.

In summary, we used 3D eye-specific models to analyze the morphology, hemodynamics, and oxygenation for ONH vasculature. Results from this study help us understand the association between the structure and function of ONH. Our model can predict tissue oxygenation from limited measurement. It can potentially lead to new biomarkers for ONH hemodynamics/oxygenation, which contribute to the glaucoma diagnosis and management. This approach may serve as a foundation for identifying novel biomarkers related to ONH hemodynamics and oxygenation. These biomarkers have the potential to improve the diagnosis and management of glaucoma.

## Acknowledgements

Supported by National Institutes of Health (R01-EY023966, R01-EY031708, R01-HD083383, P30-EY008098, T32-EY017271); Eye & Ear Foundation of Pittsburgh; Research to Prevent Blindness (unrestricted grant to UPMC Ophthalmology and Stein Innovation Award to IAS); and BrightFocus Foundation.

## Disclosures

Y. Lu, None

Y. Hua, None

R. Gill, None

A Theophanous, None

P-Y. Lee

I.A. Sigal, None

## Funding

National Institutes of Health (R01-EY023966, R01-EY031708, R01-HD083383, P30-EY008098, T32-EY017271); Eye & Ear Foundation of Pittsburgh; Research to Prevent Blindness (unrestricted grant to UPMC Ophthalmology and Stein Innovation Award to IAS); and BrightFocus Foundation.

## Notes

### Competing Interest Statement

The authors have declared no competing interest.

