## supplementary material for "Associations between structural and functional hemodynamics in the optic nerve head"

**Parameter distribution fittings.**

Table S1 summarizes the fitted distributions for all vascular structural and functional parameters across four eyes. The observed parameter distributions were fitted using different functions selected according to their shapes: quadratic for depth, peripheral distance, and blood pressure; logistic for inclination angle; exponential for tortuosity and blood flow; and skew-normal for length, blood oxygenation, and tissue oxygenation.

|  | **Fitting distribution function** | **Parameter** | **Eye1** | **Eye2** | **Eye3** | **Eye4** |
| --- | --- | --- | --- | --- | --- | --- |
| Depth(µm) | ax^2^+bx+c | a (µm^-3^) | -1.8e-8 | -2.1e-8 | -3.9e-8 | -1.7e-8 |
|  |  | b (µm^-2^) | 7.9e-6 | 1.1e-5 | 2.0e-5 | 9.4e-6 |
|  |  | c (µm^-1^) | 1.6e-3 | 1.4e-3 | 9.1e-4 | 1.4e-3 |
| Peripheral distance (µm) | ax^2^+bx+c | a (µm^-3^) | 2.1e-9 | 8.0e-10 | 2.8e-9 | 1.3e-9 |
|  |  | b (µm^-2^) | -4.9e-6 | -3.4e-6 | -5.3e-6 | -3.9e-6 |
|  |  | c (µm^-1^) | 2.7e-3 | 2.3e-3 | 2.6e-3 | 2.5e-3 |
| Inclination angle(rad) | $y=\frac{U}{1+e^{-k(x-x_{0})}}$ | U | 0.85 | 0.98 | 2.05 | 1.16 |
|  |  | k | -5.4 | -3.5 | -2.82 | -3.38 |
|  |  | X_0_ | 1.1 | 0.94 | 0.32 | 0.75 |
| Polar angle(rad) | C | C | 2/π | 2/π | 2/π | 2/π |
| Length (µm) | $\frac{1}{w\sqrt{2\pi}}e^{\frac{-{(x-x_{0})}^{2}}{2w^{2}}}[1+\mathrm{erf}(\frac{\alpha(x-x_{0})}{w\sqrt{2}})]$ | x0 (µm) | 6.3 | 12 | 13 | 13 |
|  |  | w (µm) | 47 | 34 | 50 | 37 |
|  |  | *α* | 10 | 4.2 | 6.0 | 4.3 |
| Tortuosity | $y=\lambda e^{-\lambda(x-1)}$ | $\lambda$(µm^-1^) | 9.1 | 11 | 3.8 | 11 |
| Blood flow (nl/min) | $y=\lambda e^{-\lambda(x-1)}$ | $\lambda$(min/nl) | 0.24 | 0.32 | 0.31 | 0.29 |
| Blood Pressure (mmHg) | ax^2^+bx+c | a (mmHg^-3^) | -4.4e-5 | 7.3e-5 | -2.7e-5 | 5.6e-5 |
|  |  | b (mmHg^-2^) | 2.2e-3 | -6.7e-3 | -1.1e-4 | -5.1e-3 |
|  |  | c (mmHg^-1^) | 7.9e-3 | 0.16 | 0.065 | 0.12 |
| Blood O_2_ (mmHg) | $\frac{1}{w\sqrt{2\pi}}e^{\frac{-{(x-x_{0})}^{2}}{2w^{2}}}[1+\mathrm{erf}(\frac{\alpha(x-x_{0})}{w\sqrt{2}})]$ | x0 (mmHg) | 69 | 65 | 64 | 74 |
|  |  | w (mmHg) | 6.4 | 11 | 11 | 15 |
|  |  | *α* | -4.3e-3 | -2.5e-2 | -6.6e-3 | -2.8 |
| Tissue O_2_ (mmHg) | $\frac{1}{w\sqrt{2\pi}}e^{\frac{-{(x-x_{0})}^{2}}{2w^{2}}}[1+\mathrm{erf}(\frac{\alpha(x-x_{0})}{w\sqrt{2}})]$ | x0 (mmHg) | 68 | 58 | 57 | 66 |
|  |  | w (mmHg) | 11 | 11 | 9.5 | 16 |
|  |  | *α* | -8.6 | -0.19 | -0.012 | -6.7 |

**Table S1.** The fitted coefficients for each parameter across the four eyes.

**Comparison with previous studies**

Our previous work (Lee et al., 2022) showed that ONH vessel segment length distributions are similar in shape to those reported by Jafarnejad et al. (2019) and Hahn et al. (2019), despite differences in tissue type and species. Microvascular density and tortuosity in this works are also comparable to values measured for other tissues and species (Table S2).

|  | Our work | Previous work |
| --- | --- | --- |
| Microvascular Density (mm^-2^) | 452±58 | Human brain: 328 ± 117  (Kubíková et al 2017) |
|  |  | Mouse brain: 780± 100  (Benyi et al 2017) |
| Tortuosity | 1.129±0.27 | Human retina: 1.167 ± 0.028  (Lee et al 2018) |

**Table S2**. Comparison of microvascular density and tortuosity between our work and previous studies.

**Effect of spatial autocorrelation**

The table below compares unadjusted and spatially adjusted correlations for selected parameter pairs in Eye 1. As expected, spatial adjustment reduced the magnitudes of the correlation coefficients and enlarged the p-values. The associations of depth with polar angle and tortuosity were no longer statistically significant at the 0.05 level. However, the positive association between blood flow and tissue oxygenation remained statistically significant, although its magnitude decreased from 0.467 to 0.303 (both p<1e-5).

| Parameter 1 | Parameter 2 | Unadjusted r | Spatially adjusted r | Unadjusted p | Spatially adjusted p |
| --- | --- | --- | --- | --- | --- |
| Depth | Polar angle | -0.0321 | -0.016 | 0.00283 | 0.137 |
| Depth | Inclination angle | 0.204 | 0.133 | 1e-5 | 1e-5 |
| Depth | Length | 0.0583 | 0.0234 | 1e-5 | 0.0295 |
| Depth | Tortuosity | 0.0214 | 0.00632 | 0.0466 | 0.557 |
| Peripheral distance | Polar angle | 0.0364 | 0.0261 | 0.000716 | 0.0154 |
| Peripheral distance | Inclination angle | -0.0395 | -0.0314 | 0.000236 | 0.0035 |
| Peripheral distance | Length | 0.0477 | 0.0348 | 1e-5 | 0.0012 |
| Peripheral distance | Tortuosity | 0.00132 | -0.000267 | 0.902 | 0.98 |
| Flow | Tissue O2 | 0.467 | 0.303 | 1e-5 | 1e-5 |

**Table S3**. The correlation and p value before and after address the spatial autocorrelation.
